# Scale matters: the arbuscular mycorrhiza - root hair trade-off from individual plants to communities

**DOI:** 10.64898/2026.10.05.756675

**Authors:** Elsa Matthus, Justus Hennecke, Karla M. Barfuss, Tilo Henning, Liesje Mommer, Joana Bergmann

## Abstract

1. Plant roots employ multiple strategies to acquire soil resources, including the outgrowth of root hairs and the association with arbuscular mycorrhizal fungi (AMF). These structures represent alternative pathways to access soil resources. Evidence is derived mostly from studies within single species or multi-species experiments under controlled conditions. It remains unclear whether this trade-off holds under natural conditions and whether it manifests consistently across different scales of biological organization.
2. We analysed 81 grassland species representing grasses, forbs and legumes, sampled under heterogeneous field conditions in three regions across Germany, allowing for analyses (i) across different species (interspecific), (ii) across communities (intercommunity), (iii) within plant species (intraspecific) and (iv) within root systems of individual plants (intra-individual). Fine root traits related to AMF colonization (including scoring of fungal hyphae and arbuscules) and root hair growth (including length and incidence) were quantified.
3. Comparing plant species, we observed a strong negative correlation between mycorrhizal colonization and root hair traits, confirming previous studies working under controlled conditions. The negative relationship between root hair length and mycorrhization was also evident when comparing mean trait values at the plant community scale. Among different individuals of the same species (i.e. intraspecific scale), there was no consistent pattern of trait-trait relationship. When comparing within roots of the same plant (i.e. intra-individual scale), co-occurrence of root hairs and mycorrhiza was less likely than what would be expected by chance, with forbs showing the most consistent negative pattern.
4. Our study demonstrates that the hypothesized trade-off between AMF and root hairs is a robust pattern under field conditions at the community and interspecific scale. At the intraspecific scale, no uniform trade-off pattern can be discerned, and within single root systems the negative trade-off pattern is widespread but weak. This indicates that ecological and evolutionary constraints shape trait-trait relationships at higher scale, whereas physiological limits do not restrict trait co-occurrence at lower scales.

## Introduction

Plants acquire belowground resources from the soil via their absorptive fine-root system. Two main modes of resource uptake exist: directly from soil into outermost root cells following a “do-it-yourself” strategy, or indirectly by outsourcing resource uptake to arbuscular mycorrhizal fungi (AMF;Bergmann et al., 2020; Freschet et al., 2020; Matthus et al., 2025; Smith et al., 2011). Physiologically, direct import relies on uptake mechanisms into epidermal root cells, with root hair cells being specialized epidermal cells extending up to a few millimetres into the soil (Grierson et al., 2014). Indirect uptake relies on AMF taking up resources from the soil and delivering them via their extensive hyphal network to the host plant (Martin & van der Heijden, 2024; Stewart et al., 2026). These two strategies can be viewed as functional alternatives, shaped by environmental conditions and genetic predisposition, yet they may also be constrained by anatomical and physiological limits that could render them mutually exclusive.

Both root hair outgrowth and engaging with AMF are comparable in that they are dynamic processes, can be found widespread across the plant kingdom, and increase the exploited soil volume (see Table 1 for a comparison). Both strategies also come with a carbon cost to the plant (Hawkins et al., 2023; Raven et al., 2018). Root hair construction takes place as tubular tip growth of plant epidermal cells, increasing the total root membrane surface area two to nine-fold (Holz et al., 2018; Jungk, 2001). When hosting intraradical AMF, the plant supplies carbon to feed the fungi as well as having to remodel intracellular processes to accommodate fungal structures in its cortex cells (Cargill et al., 2025; Hawkins et al., 2023). Indeed, intraradical arbuscular growth has been described with the analogy of “inward tip growth with a continually increasing number of tips”, making it a photographic negative of root hair outgrowth (Roeder et al., 2021).

**Table 1:**
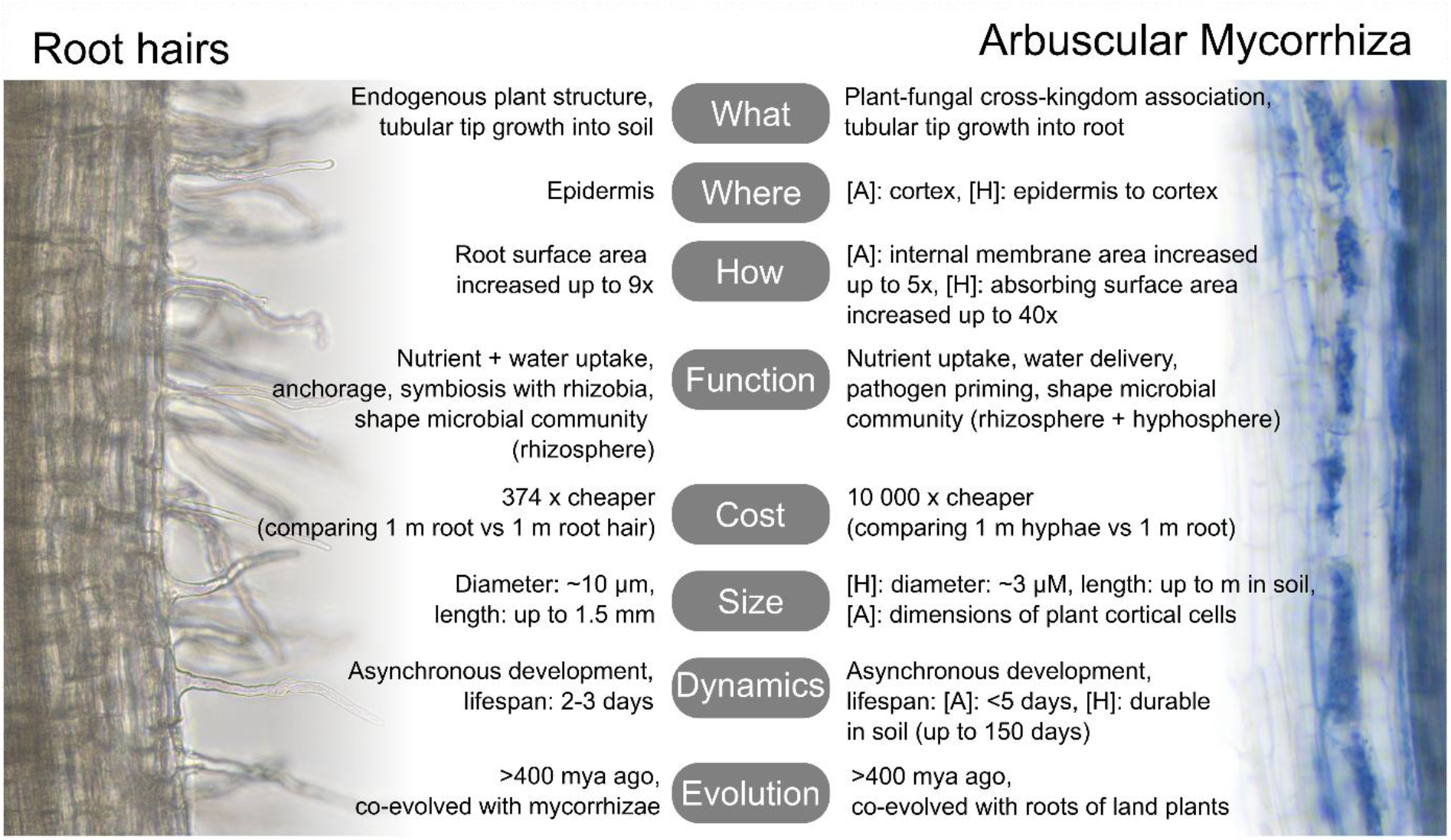
Comparison between root hair and arbuscular mycorrhiza fungal structures. Microscopic image of a field-grown root with extending root hairs (on the left), and root with arbuscular mycorrhizal fungal structures (on the right, arbuscules appear as blue cloudy structures filling cortical cells, hyphae as thin blue threads). [A] – arbuscule, [H] – hyphae. For references, see Supplementary Table S1.

It has been over 150 years that these two resource uptake strategies have been conceptually linked (Drude, 1873; Kelley, 1950). In the 1970ies, Baylis hypothesized that root morphology (including root hairs) determines to what extent plant species engage with mycorrhizal fungi, suggesting the two as alternative strategies of resource uptake (Baylis, 1970; Baylis, 1975).

Whilst numerous studies investigated either root hair *or* mycorrhizal colonization, few studies have quantified both to empirically test the hypothesized trade-off (Rongsawat et al., 2021; Zou et al., 2019). An early study across 23 orders of tropical trees and shrubs described a negative relationship between root hair outgrowth and mycorrhizal colonization (St John, 1980). A negative relationship between AMF colonization and root hair density and/or length was further reported on the intraspecific scale for the tree *Alnus glutinosa*, the grasses *Hordeum vulgare* and *Sorghum bicolor*, as well as across four monocot families (Betekhtina et al., 2023; Brown et al., 2013; Orfanoudakis et al., 2010; Sun & Tang, 2013; Wang et al., 2025). In contrast, multiple studies working on the intraspecific scale with trifoliate orange (*Pocirus trifoliata*) reported a synergistic effect of AMF inoculation on root hair outgrowth, as root hair density increased upon AMF inoculation (Hu et al., 2024; Liu et al., 2018; Wu et al., 2016; Zou et al., 2017). From these inter- and intraspecific studies, the relationship between mycorrhizal colonization and root hair outgrowth can be summarized as unclear. Recently, a study working with a large species set including over 80 grassland species reported a phylogenetically conserved trade-off pattern between root hair outgrowth and mycorrhization on the interspecific scale (Bergmann et al., 2026). The study further reported a slight negative intraspecific pattern, but replicate number per species was limited and patterns varied strongly between species (Bergmann et al., 2026).

As described above, there are first attempts to quantitatively assess the relationship between root hairs and mycorrhization on inter- and intraspecific scale. Yet, we could not find studies on the community or intra-individual scale. Whilst trait-trait relationships are often implicitly assumed to be consistent across scales of biological organization, patterns have been shown to vary (also known as the Simpson’s paradox, (Allison & Goldberg, 2002; Anderegg et al., 2018; Matthus et al., 2025). This inconsistency arises because different processes dominate at different scales:

At the interspecific scale, variation is driven by evolution (Felsenstein, 1985; Reich, 2014; Westoby et al., 2002). It displays functional biodiversity and therefore provides the largest amount of variation that nature can display. Trade-offs are typically tested at this scale, for example in root economics space and plant economics concepts (Bergmann, Weigelt, Plas, et al., 2020; Weigelt et al., 2021). At the community scale, patterns emerge from a combination of species turnover, environmental filtering, and intraspecific variation (Lepš et al., 2011; Siefert et al., 2015). We can determine community-weighted trait means as estimates of the most suited trait syndrome for a given environmental context. These are largely driven by the most dominant species in the system. Within species, at the intraspecific scale, trait variation is driven by plastic responses to local environmental conditions and different genotypes (Siefert et al., 2015). Consequently, the strength and even the direction of trait-trait relationships may weaken, disappear, or reverse as compared to other scales, as individuals adjust their trait expression to local conditions and within-species variability can be substantial (Han et al., 2025; Rodríguez-Alarcón et al., 2024). When observing one individuum only, at the intra-individual scale, there is no genetic variation. However, micro-environmental heterogeneity and epigenetics can still lead to localized phenotypic plasticity, such as “patchy” expression of traits within a single root system. Few studies have focussed on variation at this scale although it has recently gained attention (Castro Sánchez-Bermejo et al., 2025; Herrera, 2017). A trade-off at this scale would be governed by physiological constraints, either physical exclusion mechanisms or resource budgeting (i.e. having a limited number of resources to spent in a specific area of the root system).

Our understanding of the trade-off between root hairs and mycorrhization remains focussed on controlled conditions (Bergmann et al., 2026; Hu et al., 2024; Maherali, 2014; Orfanoudakis et al., 2010; Rongsawat et al., 2021; Sun & Tang, 2013; Veresoglou & Rillig, 2014). Such settings often use seeds from homogeneous populations, resulting in low genetic variation, while phenotypic variation is induced through specific treatments that inevitably select for certain subsets of traits. Given that environmental context may shape this trade-off across scales of biological organization, we investigate root hairs and mycorrhization under field conditions, using over 80 temperate grassland species including grasses, forbs and legumes. Sampling plots cover sandy to clay-rich soils across three German regions within the Biodiversity Exploratories framework (Fischer et al., 2010), and our spatially explicit intraindividual microscopic observations allow us to assess the co-occurrence of root hair and mycorrhization traits within individual root systems. With this dataset, we test whether 1) the reported interspecific trade-off can be confirmed under field conditions and whether it is consistent at 2) community, 3) intraspecific, and 4) intra-individual scale. We then synthesize our findings across all four scales, to understand the underlying drivers of this trait-trait relationship.

## Material s Methods

### Experimental design and sampling

We sampled whole plants on grassland plots (50 m x 50 m) in all three regions of the long-term Biodiversity Exploratories project (http://www.biodiversity-exploratories.de/; Fischer et al., 2010) in April and May 2021. The regions are located across Germany: in the UNESCO Biosphere Reserve Schorfheide-Chorin (SEG) in the Northeastern lowlands, the Hainich-Dün National Park in hilly central Germany (HEG) and the UNESCO Biosphere Reserve in the mountainous Southwest Schwäbische Alb (AEG). All plots have been designated grassland plots for at least 25 years prior to sampling, with different land-use intensity between the years.

Per region, we sampled nine grassland experimental plots (27 plots in total). Per plot, we sampled the ten most abundant plant species (three replicates per species and plot), as determined by aboveground vegetation analyses of previous years (Biodiversity Exploratories Information System repository: https://www.bexis.uni-jena.de, dataset ID = 31175), resulting in a mean relative aboveground vegetation coverage of 84 % per plot (minimum: 54 %, maximum: 99 %). In total, we sampled 81 unique species, including 53 non-leguminous forbs (from here on: forbs), 5 legumes and 23 grasses. Whole plants, including surrounding bulk soil, were dug out, stored cool in the field, and later frozen at -20 °C. In the laboratory, samples were thawed, and soil gently washed away from the roots using tab water.

## Root trait measurements

Fine roots (< 2 mm in diameter) of first and second order roots were considered for the reported root traits. For measurements of root hair length, a subsample of the washed root system was stored in ROTI®Histofix ECO Plus (Carl Roth, Germany), which was later exchanged for 70 % ethanol and stored at 4 °C. A representative subsample was mounted on a microscopic slide (76 x 26 x 1 mm, Epredia, Netherlands) in a drop of water and covered with a cover glass (60 x 24 x 0.13-0.16 mm, Epredia, Netherlands). Under a light microscope (Leica DM750, Germany, with LEICA DMC6200 camera and 40 x magnification), first to second order root tips were imaged, taking 3-5 images per subsample to analyse 5-6 mm of root length in total. Each image was overlaid with a grid, dividing the image into 1mm^2^ quadrants. Per quadrant, the longest root hair cell on each side of the root was measured using the segmented line tool in open-source software ImageJ (Abràmoff et al., 2004).This yielded at least 12 root hair length measurements per sample. Root hair length could not be determined on intra-individual scale, as preparation of stained root samples (see below) does not allow for unbiased length measurements.

Remaining root tissue was oven-dried (3 days, 60 °C). For determination of mycorrhizal colonization levels and root hair incidence, the dried root sample was cut into approximately 1 cm long sections, of which a subsample was used for subsequent staining of fungal structures (Barto et al., 2010; Vierheilig et al., 1998). In short, root sections were wrapped in lace paper (Kao Germany GmbH, Germany) and secured in staining cassettes (Kartell Spa, Italy). The samples were submerged in 10 % (v/v) KOH solution, in a water bath at 80°C for 15 min. After rinsing once with distilled water, and once with 5 % (v/v) acetic acid (Merck, Germany), samples were stained with an ink-acetic mixture (10 % Shaeffer black ink, A.T. Cross Company, USA; in 5 % acetic acid, Merck, Germany), at 80 °C for 15 min. Samples were then rinsed in distilled water, covered in lactoglycerol (1:1:1 mixture of glycerol, lactic acid and deionized water; Carl Roth, Germany) and left to stand for at least 30 minutes and up to 24 hours before mounting root sections on glass slides (76 x 26 x 1 mm, Epredia, Netherlands) using a few drops of lactoglycerol as a transparent mounting medium, covered with a cover slip (60 x 24 x 0.13-0.16 mm, Epredia, Netherlands) and sealed with colourful nail polish. Prepared slides were stored at 4°C and scored under the microscope within two weeks.

The degree of mycorrhizal colonization of the roots as well as root hair incidence was evaluated using the Magnified Intersection Method (McGonigle et al., 1990) on a light microscope as described above at 200 x magnification. For each sample, we aimed to score at least 50 and ideally 100 intersects, each intersect corresponding to one observation when the crosshair of the microscope overlapped with the root sample when moving the field of view across the sample. For each intersect, the presence of mycorrhizal structures (arbuscules, vesicles, coils, and hyphae), as well as root hair presence, was scored in binary format (1 = yes, 0 = no, also see Figure 4A for schematic) in an excel spreadsheet inspired by the AMScorer (Jarratt-Barnham et al., 2024). This allowed us to record information about spatial co-localization of AMF structures and root hairs (at 200 x magnification, the field of view of the microscope covered an area of approximately 0.785 mm^2^). Both root hair incidence and mycorrhizal colonization traits can be calculated as percentages of total observations per sample. Throughout this manuscript we will be reporting “AMF colonization” as a measure for *any* intraradical fungal structure observed within the root (fungal hyphae, arbuscule, vesicle or coil), as compared to “arbuscular colonization” which quantifies observation in which arbuscules had to be present. In total, 796 samples were scored for mycorrhizal colonization and root hair incidence. Root hair length was measured on 724 samples, as 72 samples had too little sample tissue for root hair length measurements.

## Statistical analyses

All analyses were conducted in R (version 4.5.0). To quantify trait–trait relationships between root hair traits and AMF colonization traits, standardized major axis (SMA) regression models were fitted using the sma() function (smatr package, Warton et al., 2012) at three biological scales: (i) interspecific scale, using species mean trait values across all plots and regions (n = 79-81 species), (ii) community scale, using plot-specific community-weighted means (CWMs, weighted by relative aboveground species cover based on cover data from springs of four years (2018-2021), n = 27 communities), and (iii) intraspecific scale, using trait observations within species (filtering for species that were sampled in at least 6 replicates, n = 32-41 species, followed by a meta-regression using the rma.uni() function (metafor package, Viechtbauer, 2010) across all individual species SMA slopes, reporting a pooled SMA slope. On intraspecific scale, I^2^ was calculated as a measure for the proportion of total variation in effect sizes (in this case: intraspecific SMA slopes across species) that is due to true between-species heterogeneity rather than within-species sampling error. SMA analysis is particularly useful to describe trait-trait relationship if both traits have measurement error and no trait is strictly dependent. Pearson correlations were reported to further test if traits co-vary.

The analysis of the intra-individual observations of binary presence-absence data from repeated microscopic observations within individual roots required a different approach. For each individual root sample, we used bidirectional generalized linear mixed-effects models (binomial error distribution and logit link using glmer() function (lme4 package, Bates et al., 2015), modelling AMF/arbuscule occurrence as a function of root hair presence, and *vice versa*, with random intercepts and random slopes for individuals nested within species to account for repeated observations and hierarchical structure. Additionally, the same model was run including plant functional type as an interacting predictor. As a second approach to understand co-occurrence patterns, we calculated the observed frequency of co-occurrence of AMF/arbuscules and root hairs and the expected frequency under independence, following the formula:

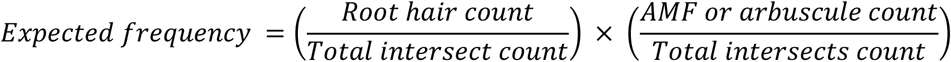

and using their difference (D) as an index of local exclusion. Within-ID permutation test (2000 permutations) was used to evaluate whether co-occurrence within individuals was lower than expected by chance, followed by Benjamini-Hochberg correction to adjust for multiple testing across individuals. A one-sided Wilcoxon signed rank test was used to assess whether D values across individuals followed a significant trend.

To test how plant functional type modified the relationship between traits, different approaches were needed as data structure differed between scales. On interspecific scale, grouped SMA regressions were fitted and tested for whether slopes differed between forbs and grasses. On community scale, the proportion of grasses versus forbs and legumes (based on their aboveground cover per plot) was calculated, and (as SMA cannot handle continuous variables) linear models with this proportion variable included as interaction term were fitted. On intraspecific scale, functional-group effects were assessed using moderator meta-regressions, in which plant functional group was included as a predictor of species-specific trait–trait slope estimates. On the intra-individual level, functional-group effects were assessed by including plant functional group as interaction term into the mixed model.

To visualize trait expression along the phylogenetic tree of species relatedness, we pruned the DaPhnE phylogeny (Durka & Michalski, 2012) to our species set using drop.tip() (ape package, Paradis et al., 2004) and plotted via phylo.heatmap() function (phytools package, (Revell, 2024).

## Results

### Comparing species, root hair outgrowth and mycorrhizal colonization are strongly negatively related

To assess if an interspecific trade-off between root hair and mycorrhization traits can be found under field conditions, we first tested the trait-trait relationships on interspecific scale (Figure 1). Species with higher mycorrhizal colonization showed significantly lower levels of root hair incidence (Figure 1A, 1C), as well as shorter root hairs (Figure 1B, 1D; with SMA slopes significantly differing from 0 and Pearson *p* < 0.01 in all four cases). This significantly negative relationship was observed for total AMF colonization (including all fungal structures such as hyphae, arbuscules and vesicles, Figure 1A, 1B) as well as for arbuscule only colonization (Figure 1C, 1D). Considering the rate of trait change, a SMA slope of -2.21 (Figure 1C) indicates that for every 10 % increase in arbuscule colonization, root hair incidence is predicted to drop by approximately 22 %.

**Figure 1:**
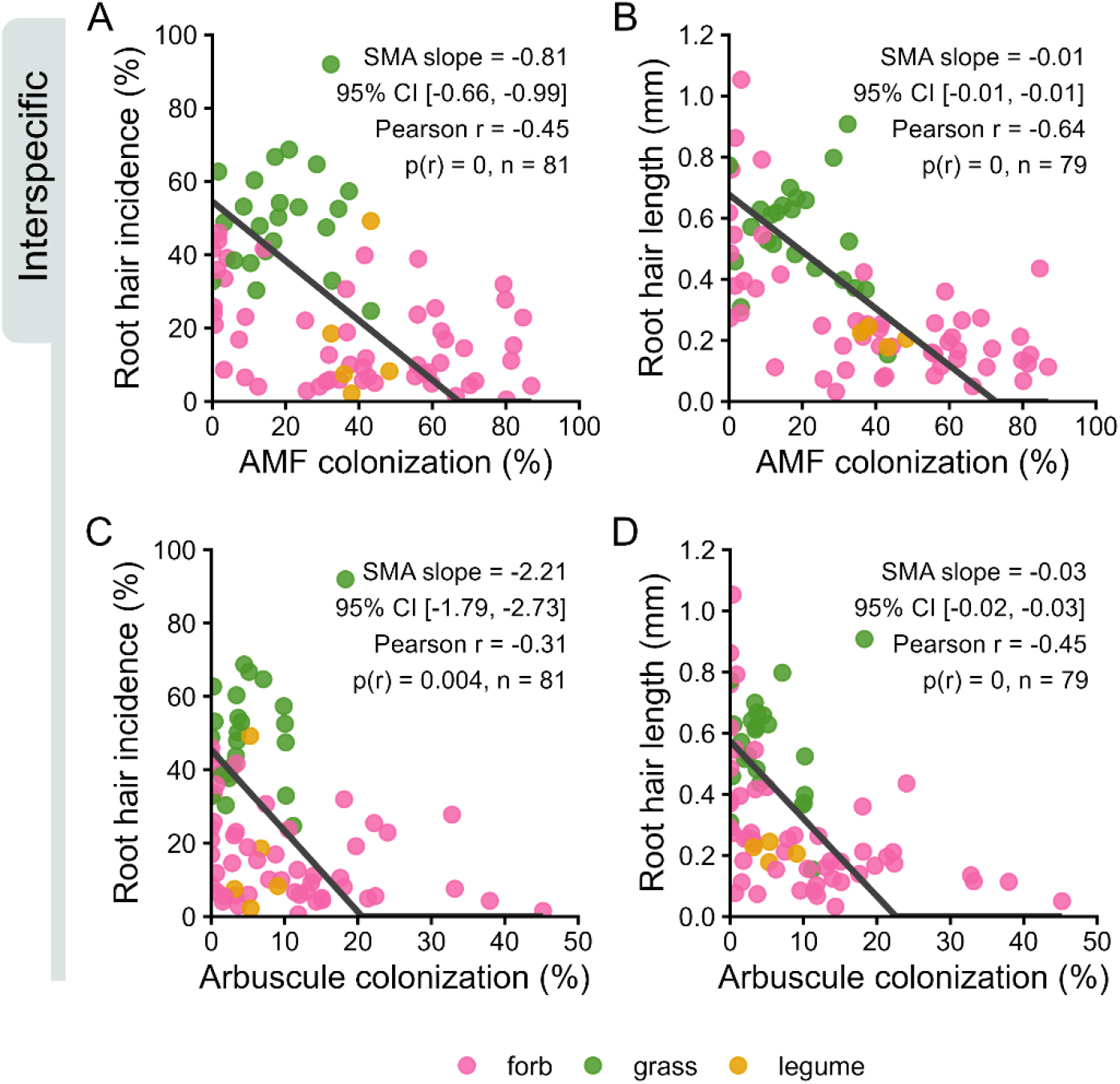
Interspecific relationship of root hair traits and mycorrhizal colonization in grassland species. Species mean trait values of root hair incidence (A, C; n = 81 species) and root hair length (B, D; n = 79 species) were plotted against total AMF colonization (A, B; including all intraradical mycorrhizal structures such as hyphae, arbuscules, vesicles, coils) and arbuscular colonization (C,D; including only arbuscules). Each point corresponds to a species mean and is coloured according to the plant functional type. Black line denotes the standardized major axis (SMA) regression slope. AMF – arbuscular mycorrhizal fungi.

This negative relationship between the two strategies was further apparent when clustering trait expression levels according to a phylogenetic tree of species relatedness: species with high values of mycorrhizal colonization levels tended to have fewer and shorter root hairs, and *vice versa* (see Supplementary Figure S1). The pattern closely replicated what had been observed in a previous study working with a different set of grassland species under controlled conditions (Bergmann et al., 2026).

We noticed that plant functional types differed in their trait expression. Grasses carried many and long root hairs and showed lower levels of AMF colonization, whilst forbs and legumes had fewer, shorter root hairs and higher levels of AMF colonization (Supplementary Figure S2). We therefore tested if the overall negative interspecific relationship between root hair outgrowth and mycorrhizal structures could equally be observed within each functional group. In all four trait-trait relationships, testing for common slopes yielded significant differences between plant functional types (Supplementary Table S2). As the number of legume species were too low to infer robust patterns (n = 5), we concentrated on forbs and grasses (n = 53, n = 23, respectively). Forbs replicated the significantly negative relationship observed when comparing across all species and plant functional types (SMA slopes different from 0, Pearson *p* < 0.01, Supplementary Figure S3). On the contrary, in grasses, the trait-trait relationships were inconsistent (as SMA slopes were significant, *i.e.* different from 0, but Pearson *p* > 0.05, Supplementary Figure S4).

Whilst forbs were in general characterized by shorter root hairs and high levels of mycorrhizal colonization, we noticed a cluster of species from the *Caryophyllaceae* family (*Stellaria media, Cerastium holosteoides, Cerastium glomeratum*) within the forbs that showed long root hairs and low-to-no mycorrhization (Supplementary Figure S1). In the case of *Cerastium holosteoides*, the average root hair length was 1.05 mm, which represented the longest root hairs of all species assessed in this study.

Taken together, root hair traits and mycorrhizal colonization structures were negatively related when comparing across species. Considering the plant functional types, forbs but not grasses followed the significantly negative pattern observed across all species.

### Comparing communities, root hair length is negatively related to mycorrhizal colonization

We next assessed trait relationships at the community scale, using community weighted mean (CWM) trait values. We detected a negative trend in all four trait-trait relationships (Figure 2). Root hair incidence showed a weakly negative but inconsistent relationship with AMF colonization and arbuscule colonization (SMA slopes differed from 0 but Pearson *p* was not significant; Figure 2A, 2C, respectively). Root hair length showed significant negative relationship with both AMF and arbuscule colonization (Figure 2B, 2D, SMA slopes differ from 0, Pearson *p* ≤ 0.001). For example, an increase in 10 % of AMF colonization is predicted to be linked to a 0.1 mm decrease in root hair length (Figure 2B).

**Figure 2:**
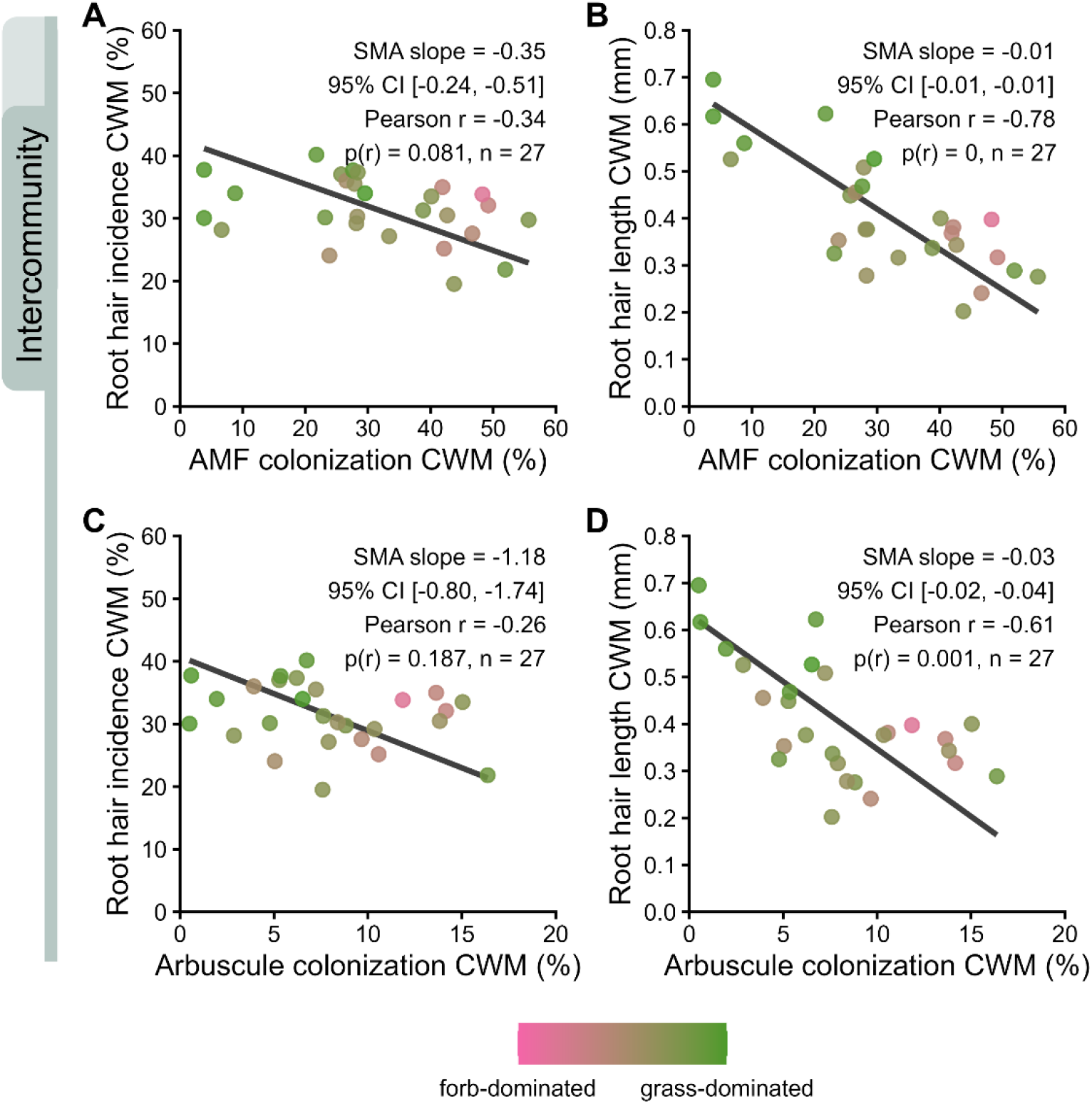
Intercommunity trait-trait relationships between root hair outgrowth and mycorrhizal colonization. Community-weighted mean (CWM) traits (weighted by aboveground plant cover) of root hair incidence (A, C) and root hair length (B, D) were regressed against AMF colonization (A,B; including all intraradical mycorrhizal structures such as hyphae, arbuscules, vesicles, coils) and arbuscular colonization (C, D). Each data point represents one community (n = 27) and is shaded according to the proportion of grasses (green) or forbs (pink), calculated based on aboveground cover per plot. Black line denotes standardized major axis (SMA) regression slope. AMF – arbuscular mycorrhizal fungi.

We observed that forb-dominated communities clustered at a specific end of the trade-off, *i.e.* at low root hair and high mycorrhization values. To assess whether plant functional type community composition influenced the trait-trait relationships, we fitted supplementary linear models including the interaction between plant functional type cover proportion and the predictor trait. Two models involving root hair length showed significant interactions (RHL ∼ AMF, RHL ∼ Arbuscules), but equivalent reverse-direction models did not support these effects (AMF ∼ RHL, Arbuscules ∼ RHL, Supplementary Table S3). We therefore concluded that interaction terms were generally not supported, indicating that the observed trait-trait relationships were largely consistent across the grass-forb composition gradient.

Collectively, results at community scale showed that high levels of mycorrhization were linked to shorter root hairs. The relationship between root hair incidence and mycorrhization showed a negative but less robust trend. As these relationships were generally consistent across grass- and forb-dominated communities, they appear to reflect a general trade-off between root morphological investment and mycorrhizal symbiosis rather than differences in plant community composition due to plant functional type dominance alone.

### Within species, root hair outgrowth and mycorrhizal colonization show varied trait-trait patterns

When comparing intraspecific relationships of root hair outgrowth and mycorrhizal colonization, the patterns were highly variable (Figure 3). Both trade-off patterns (negative slopes) as well as synergistic effects (positive slopes) could be observed within species (Figure 3, also see Supplementary Figure S5 for detailed forest plots). Heterogeneity within species was extremely high in all four trait-trait combinations (I² = 97-99 %), with species-specific slopes varying substantially in both magnitude and direction. This indicates pronounced species-specific differences in intraspecific trait coordination. When summarizing the individual SMA slopes into a pooled slope per trait-trait combination, these pooled slopes were in all cases close to 0 and mostly non-significant. When testing if plant functional types could explain the varied patterns, no functional type showed a significiant effect in any of the trait-trait relationships. Taken together, we found no clear evidence for a trade-off between mycorrhizal colonization and root hairs within species.

**Figure 3:**
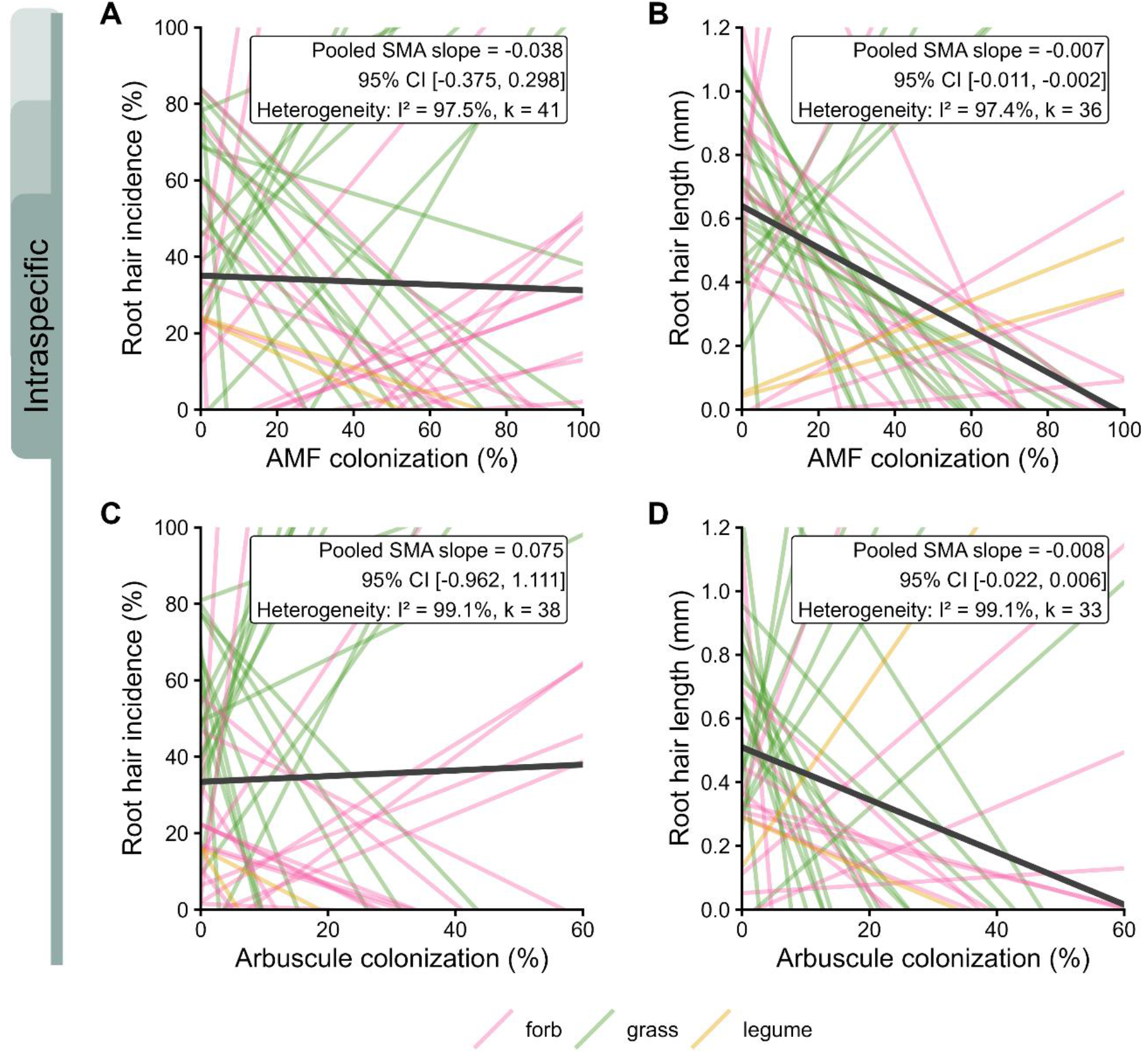
Intraspecific trait-trait relationships between root hair outgrowth and mycorrhizal colonziation. Standardized major axis (SMA) regressions were fitted per species (thin lines, coloured by plant functional types: forbs (purple), grasses (pink), legumes (orange)) to assess the relationship of root hair incidence (A, C) and root hair length (B, D) to AMF colonization (A, B) and arbuscule colonization (B, D). The dark solid line denotes the pooled SMA slope across all tested species (k = 33 – 41). Please note that whilst slopes in B and D appear steep, steepness is driven by axis scaling rather than effect size and is a visual artefact. AMF – arbuscular mycorrhizal fungi.

### Within individual root systems, root hairs and mycorrhizal structures are less likely to co-occur

At the intra-individual scale, we kept track of the spatial co-occurrence scored of root hairs and mycorrhizal structures whilst scoring (Figure 4A, see Material & Methods for details). We recorded over 67 000 individual microscopic observations in total and noticed that root systems are vast spaces: In about half of the microscopic observations, roots carried neither root hairs nor mycorrhizal structures (Figure 4B, C). In about a third of the observations, only root hairs or only mycorrhizal structures could be observed (Figure 4B). Co-occurrence was low, with only 7 % showing both root hairs and an AMF structure (Figure 4B), and only 2 % showing root hairs and arbuscules to co-occur (Figure 4C).

**Figure 4:**
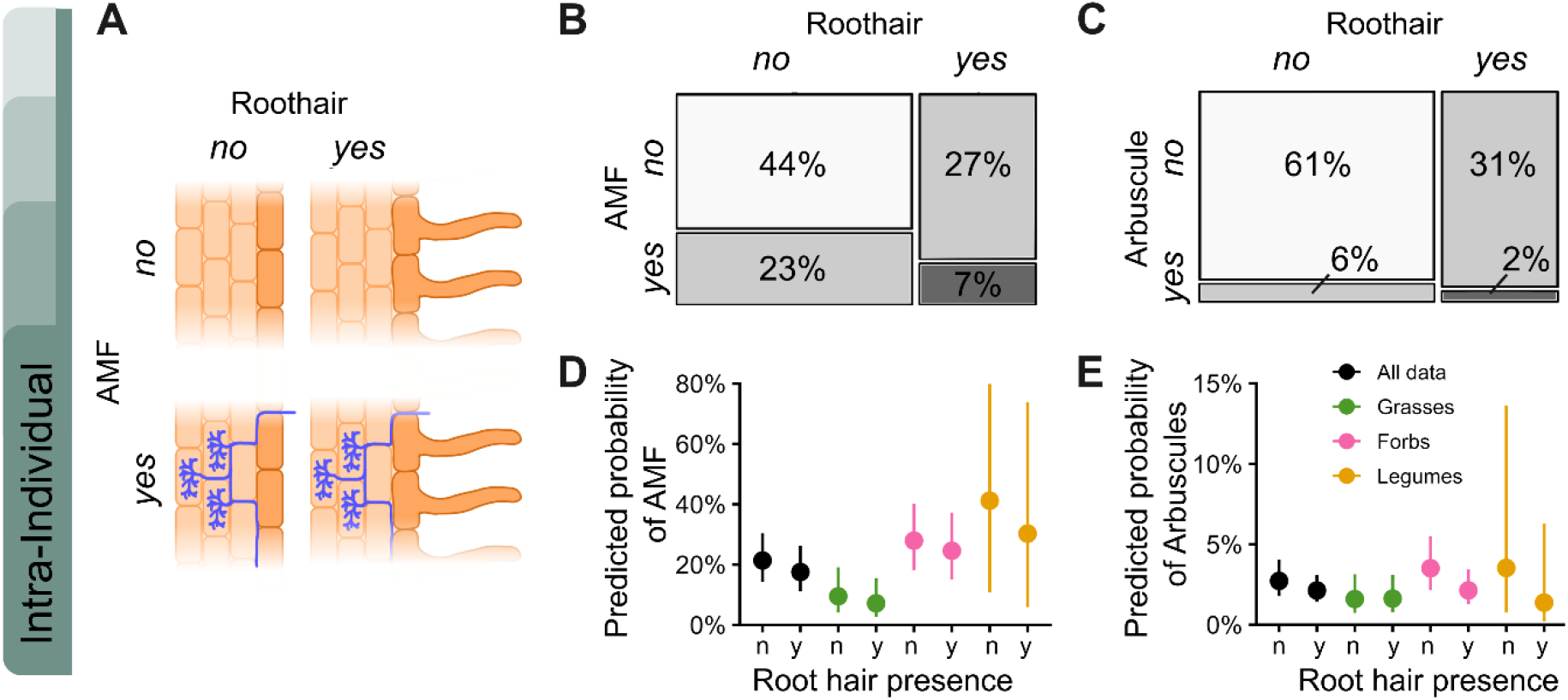
At the intra-individual scale, root hairs and mycorrhizal structures co-occur less than what would be expected by chance. (A) Schematic depicts our scoring approach of the co-occurrence of root hair (yes/no) and blue AMF structures (yes/no for hyphae, arbuscules, vesicles, coils) at each microscopic observations within root individual root system. Mosaic plots depict proportions of in total 67161 unique microscopic observations that carried (B) root hairs and/or AMF structures, (C) root hairs and/or arbuscules. Predicted probabilities of (D) AMF occurrence or (E) arbuscule occurrence depending on root hair presence (y – yes, n – no), across all observations (black) and split by plant functional type. AMF – arbuscular mycorrhizal fungi.

Initial logistic regression models across all individual observations suggested a significantly negative association, with root hair presence reducing the probability of AMF occurrence by approximately 4 to 6 % (Figure 4D, black dots) and *vice versa*, AMF presence reducing the probability of root hair occurrence by 3 to 5 % (Supplementary Figure S6A). Similar patterns were observed when considering arbuscules: root hair presence reduced the probability of arbuscule occurrence by 2 to 4 % (Figure 4E, black dots), whereas conversely, arbuscule occurrence reduced the probability of root hair occurrence by approximately 6 to 10 % (Supplementary Figure S6B, Supplementary Table S4).

Co-occurrence analyses across all individual root systems supported these results, as the distribution of D values was slightly skewed toward negative values (Wilcoxon *p* < 0.001), indicative of root hair and mycorrhizal structures co-occurring less than what would be expected by chance (Supplementary Figure S7). However, only a small fraction of individual root systems showed exclusion patterns strong enough to remain significant after correcting for multiple testing (10 out of 652 root systems for AMF colonization, and 1 out of 498 for arbuscule colonization; Supplementary Figure S8). This indicates that negative co-occurrence patterns are weak and widespread rather than strong within individual roots.

We next assessed if the negative pattern across all observations would be modulated by plant functional type. Considering the relationship between AMF occurrence and root hairs, the pattern observed across all observations was replicated in all plant functional types, with reciprocal models yielding consistent results (Figure 4D, Supplementary Figure S6). Interestingly, this was not the case when considering root hair and arbuscule occurrence. Here, grasses deviated from the observed global pattern, as they did not show a drop in predicted probability of root arbuscule occurrence depending on root hair presence and *vice versa* (Figure 4E).

Together, these results indicate that at the level of the individual root system, root hairs and mycorrhizal structures are less likely to co-occur than what would be expected by chance. However, in the case of the functional unit of mycorrhizal symbiosis, the arbuscules, this trade-off emerged primarily in the plant functional type of forbs and was absent in grasses.

## Discussion

In this study, we tested if the trade-off between root hair formation and colonization with arbuscular mycorrhizal fungi (AMF), previously reported under controlled growth conditions (Bergmann et al., 2026), would hold under field conditions and across different scales of biological organization. By sampling 81 grassland species in 27 plots across Germany, we found that the relationship between root hairs and mycorrhizal structures was strongly scale-dependent. Whilst a trade-off was observed at the interspecific and partly at the community scale, it was absent at the intraspecific scale, where both positive as well as negative trait-trait coordination occurred and was not explained by plant functional type. At the intra-individual scale, co-occurrence of root hairs and mycorrhizal structures was less likely than what would be expected by chance.

Previous work investigating root hairs and AMF as coordinated or divergent strategies of resource uptake has provided valuable insights into species strategies under common growing conditions, mostly in greenhouse settings (Bergmann et al., 2026; Brown et al., 2013; Orfanoudakis et al., 2010; X. G. Sun & Tang, 2013). Field conditions, in contrast, encompass much wider environmental variation, potentially greater genetic variation, older plant populations and more complex soil microbial communities. Broadening these drivers when moving from controlled to field conditions, as reported here, resulted in qualitatively and quantitatively similar interspecific trait-trait relationships (compared to e.g. Bergmann et al. 2026). This supports the prevailing paradigm (Baylis, 1970) that different plant species indeed use root hairs and AMF as alternative strategies of resource uptake.

Trait-trait relationships are most commonly studied at the interspecific scale (e.g. Bergmann, Weigelt, Van Der Plas, et al., 2020; Matthus et al., 2025; Reich, 2014; Weigelt et al., 2021) which is also where we found the clearest trade-off. At this scale, trait differences reflect evolutionary history and species-level ecological strategies (Bergmann, Weigelt, Plas, et al., 2020; Westoby et al., 2002). Although our species pool was restricted to temperate grassland species, it covered considerable phylogenetic diversity found in these systems, allowing large ecological and life-history differences among species and lineages to emerge (Anderegg et al., 2018). Specifically, the phylogenetically conserved shift from root hairs to AMF in Asteraceae and the opposite trend in Poaceae contributed to the strong interspecific trade-off (also see Bergmann et al., 2026).

The longest root hairs were found in species belonging to the Caryophyllales, an order for which evidence is accumulating that they have lost the capacity to engage with mycorrhizal fungi (Lambers & Teste, 2013; Montero et al., 2025). Caryophyllales have instead been described to engage with other fungal endophytes (Giesemann et al., 2020), and their intraradically growing hyphae may resemble AMF structures, incorrectly suggesting low levels of “AMF colonization” in these field-grown plants. The absence of arbuscules (as a very specific read-out of AMF colonization) in these species in our study, however, agrees with their loss of functional mycorrhizal capacity. We could not find reports detailing their extreme root hair phenotype or explaining its genetic basis. However, it resembles what has been described for the Proteaceae where loss of the capacity to engage with mycorrhizal fungi coincides with pronounced root hair outgrowth (Lambers et al., 2015).

Community-scale trait expression is the result of many drivers. As we calculated community-weighted mean traits from plot-specific species traits, our estimates incorporate intraspecific trait variation rather than relying on fixed species means (Lepš et al., 2011). However, the absence of a consistent intraspecific trait-trait pattern suggests that within-species shifts are unlikely to be the main driver of the community scale pattern. Instead, species turnover and environmental filtering may translate the interspecific trade-off into community-scale trait composition. This is consistent with previous work showing that community-weighted mean traits can shift through both intraspecific trait variation and species turnover (Lepš et al., 2011), and that these components may differ in strength or even oppose each other along environmental gradients (Bruelheide et al., 2018; Kichenin et al., 2013).

The difference between the two root hair traits, length and incidence, is informative in this context. The interspecific trade-off between root hair incidence and mycorrhization was absent at the community scale, suggesting that species position along this trade-off is not strongly associated with community composition across our sites. In contrast, the trade-off between root hair length and mycorrhization persisted at the community scale, suggesting that relative investment in mycorrhizal association versus root hair extension is linked to how species are filtered into, and weighted within, local communities.

Trait-trait relationships at the intraspecific scale result from potentially very different drivers than interspecific relationships. Although genetic variation exists within a species, evolutionary history becomes less important, while environmental plasticity becomes more prominent (Anderegg et al., 2018). The absence of a general intraspecific trade-off in our study therefore suggests that environmental variation did not lead to a coordinated shift in resource uptake strategies within species.

This contrasts with studies that found an intraspecific trade-off along experimental gradients of symbiont availability (Ma et al., 2021; Orfanoudakis et al., 2010; Sun & Tang, 2013) or nutrient (especially phosphate) availability (Brown et al., 2013; Wang et al., 2025). These studies, however, each investigated a single species, with a strong focus on Poaceae, such as sorghum, barley and maize, along steep biotic or abiotic gradients. Our dataset covered a larger number of species, yet with only limited coverage of most species along large-scale environmental gradients: of 81 species sampled, 40 were represented by fewer than six plant individuals. Varying sample numbers have resulted from our experimental design that focussed on sampling the ten most abundant species per plot.

Nevertheless, even among species with higher replicate number, we observed trade-off, neutral and synergistic trait relationships that could not be explained by plant functional type. For example, grasses showed both positive as well as negative trait-trait relationships (Supplementary Figure S5), indicating that the trade-offs reported for individual grass species (Ma et al., 2021; Orfanoudakis et al., 2010; Sun & Tang, 2013; Wang et al., 2025) only capture part of the variation in plasticity present in larger species sets. This is particularly relevant for agricultural applications and plant breeding, where identifying optimal phenotypes increasingly includes belowground traits such as root hairs and mycorrhizal responsiveness (Grondin et al., 2024; Tsang et al., 2024). Our results suggest that coordination between these two acquisition strategies is species-specific and context-dependent, rather than following a general intraspecific trade-off. This underlines the results presented in Bergmann et al (2026) who found weak support for an intraspecific trade-off despite strong interspecific differences. Improving plant performance in specific environments may require a more nuanced understanding of how root hair traits and mycorrhizal colonization are coordinated within individual crop species or genotypes.

At the intra-individual scale, negative co-occurrence between root hairs and AMF was widespread across all root systems (= considering all data) but weak within individual root systems (= considering each root system separately and accounting for multiple testing). Phrased differently: given the fact that root hairs and mycorrhizal structures *could* spatially co-occur, a strict physiological exclusion mechanism at this scale seems unlikely. The trade-off at higher scales is therefore unlikely to arise simply from local exclusion at lower scales.

Importantly, these patterns do not resolve whether the relationships between root hair outgrowth and mycorrhization is directional – whether root hair outgrowth impedes AMF colonization or, vice versa, AMF colonization inhibits root hair outgrowth. Both mechanisms are biologically plausible. Initial AMF penetration has been reported to rely on signalling cascades in atrichoblasts, i.e. epidermal cells that have not proliferated into root hairs (Chabaud, 2011; Sun, 2015). Conversely, root hair outgrowth can depend on positional cues from underlying cell files (Dolan, 1993; Lee, 2024), raising the possibility that arbuscule growth in cortical cells could influence cortex-epidermal organisation, although to our knowledge this has not been tested. Because neither trait could therefore be clearly treated as dependent on the other, we used statistical approaches that account for this symmetry when relating the two traits, including standardized major axis regression or bi-directional models (Warton, 2006).

Our spatial analysis also has an important constraint: co-occurrence was determined within the microscope field of view, which does not allow cellular resolution. However, our spatial resolution captured microscale heterogeneity, including patchy nutrient hot spots or a localized pathogen response, as well as variation arising from tissue heterogeneity. Each individual root system can thus be viewed as a complex landscape in which plant roots and mycorrhizal fungi meet.

Taken together, our analysis across biological scales shows that interspecific trade-offs between root hairs and mycorrhization reported from greenhouse studies are also present in natural communities, but do not propagate consistently across lower scales of organization. Evolutionary history appears to be a dominant contributor to interspecific trait expression and trade-offs, whereas within-species plasticity produced less consistent patterns under field conditions. We conclude that the relationship between root hair outgrowth and mycorrhizal engagement is not a single biological law, but a scale-dependent pattern generated by different mechanisms at different levels of organization.

## Acknowledgements

EM, KMB and JB thank the DFG for funding this research (HAIRphae project, SPP 1374, 432975993), LM and JH acknowledge funding of the ERC-AdG grant number 101141742 – DIGDEEP. We further thank Julian Ahlborn for his expertise during plant identification in the field, and our interns Katia Pfister and Helga Hoppe for their help during sample preparation at the Paulinenaue Experimental Infrastructure Platform. We are very grateful for fruitful discussions within the Sustainable Grassland Systems group.

We thank the managers of the three Exploratories, Julia Bass, Miriam Teuscher, Franca Marian and all former managers for their work in maintaining the plot and project infrastructure; Victoria Grießmeier for giving support through the central office, Cornelia Fürstenau and Andreas Ostrowski for managing the central data base, and Markus Fischer, Eduard Linsenmair, Dominik Hessenmöller, Ingo Schöning, François Buscot, Ernst-Detlef Schulze, Wolfgang W. Weisser and the late Elisabeth Kalko and Daniel Prati for their role in setting up the Biodiversity Exploratories project. We thank the administration of the Hainich national park, the UNESCO Biosphere Reserve Swabian Alb and the UNESCO Biosphere Reserve Schorfheide-Chorin as well as all landowners for the excellent collaboration.

## Author contributions

Joana Bergmann conceived and designed the original experiment; Elsa Matthus, Karla Barfuss, Tilo Henning, Justus Hennecke and Joana Bergmann collected the data; Elsa Matthus and Justus Hennecke analysed the data; Elsa Matthus led the writing of the manuscript and produced all figures. All authors contributed critically to the drafts and gave final approval for publication.

## Data availability statement

All data available from the Biodiversity Exploratories Information System repository: https://www.bexis.uni-jena.de. Dataset IDs = 31827; 32391.

## Supplementary Information

**Supplementary Table S1:** Reference list corresponding to Table 1.

|  | **Root hair statement** | **Reference** |
| --- | --- | --- |
| What | Endogenous plant structure; tubular tip growth into soil | (Datta et al., 2011; Grierson et al., 2014) |
| Where | Epidermis | (Datta et al., 2011; Grierson et al., 2014) |
| How | Root surface area increased up to 9x | (Holz et al., 2018; Jungk, 2001; Ruiz et al., 2020) |
| Function | Nutrient + water uptake; anchorage; symbiosis with rhizobia; shaping microbial community (rhizosphere) | (Datta et al., 2011; Haling et al., 2013; Oldroyd, 2013; Robertson-Albertyn et al., 2017) |
| Cost | 374x cheaper (comparing 1 m root vs 1 m root hair) | (Raven et al., 2018) |
| Size | Diameter: ~10 µm; length: up to 1.5 mm | (Datta et al., 2011; Grierson et al., 2014) |
| Dynamics | Asynchronous development; lifespan: 2-3 days | (Datta et al., 2011; Feng et al., 2026; Jungk, 2001) |
| Evolution | >400 mya ago, co-evolved with mycorrhizae | (Brundrett, 2002; Jones & Dolan, 2012; Kenrick & Strullu-Derrien, 2014) |
|  | **Arbuscular mycorrhiza statement** | **Reference** |
| What | Plant-fungal cross-kingdom association; tubular tip growth into root | (Parniske, 2008) |
| Where | A: cortex; H: epidermis to cortex | (Cargill et al., 2025) |
| How | A: Increase of internal membrane area (up to 5x), H: increase of absorbing surface area (up to 40x) | (Pepe et al., 2018; Roeder et al., 2021) |
| Function | Nutrient uptake; water delivery; pathogen priming; shape microbial community (rhizosphere + hyphosphere) | (Frew, 2025; Kakouridis et al., 2022; Qin et al., 2022; Wang et al., 2023) |
| Cost | 10 000x cheaper (comparing 1 m hyphae vs 1 m root) | (Raven et al., 2018) |
| Size | H: diameter ~3 µm; length up to metres in soil; A: dimensions of plant cortical cells | (Cargill et al., 2025; McGaley et al., 2026; Stewart et al., 2026) |
| Dynamics | Asynchronous development; lifespan: A <5 days within cortex cell; H: durable in soil (up to 150 days) | (McGaley et al., 2026; Pepe et al., 2018; Stewart et al., 2026) |
| Evolution | >400 Mya ago; co-evolved with roots of land plants | (Brundrett, 2002; Christine et al., 2018; Taylor et al., 1995) |

**Figure S1:**
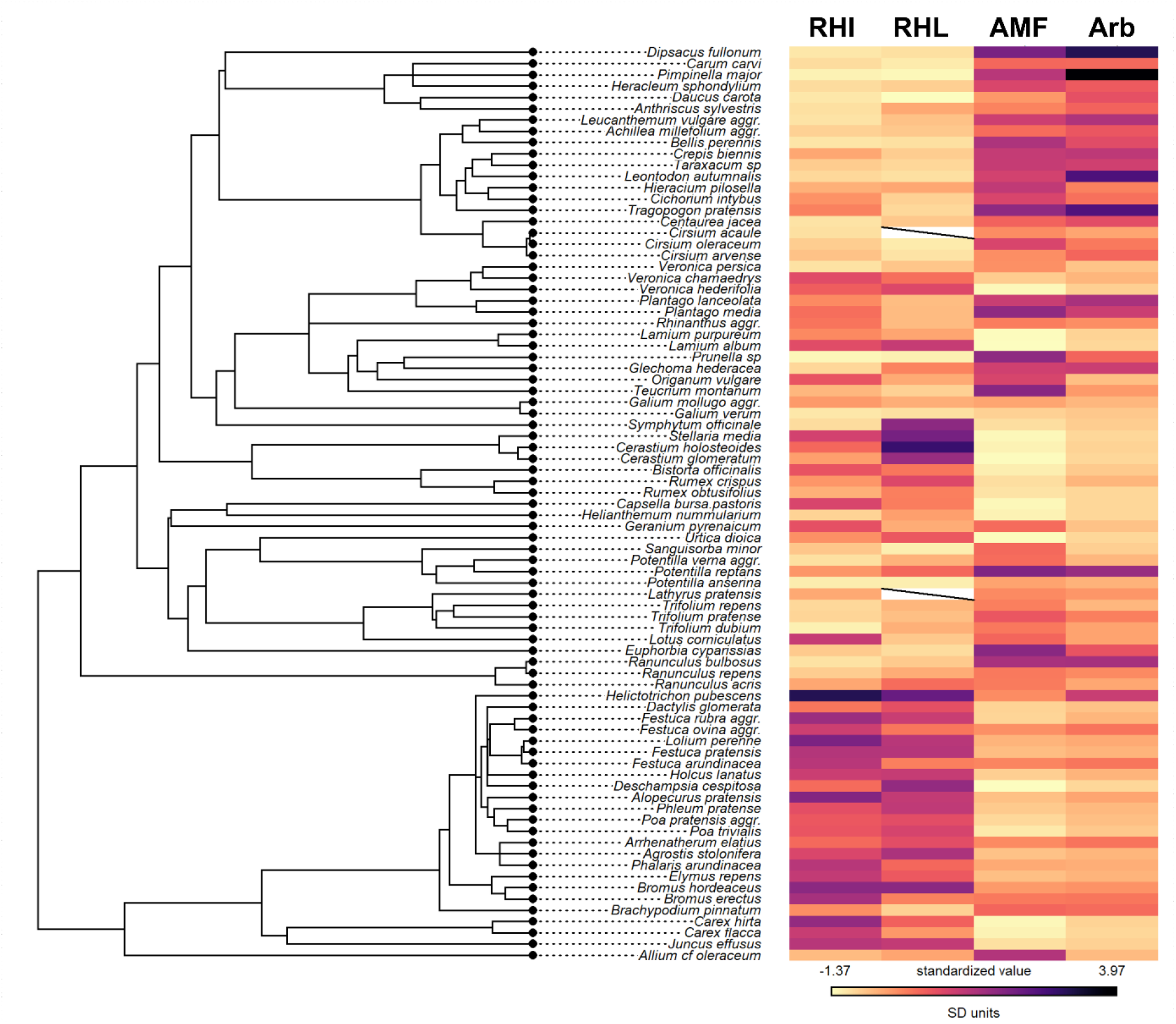
Phylogenetic tree of grassland species included in study (left) and heat map of their trait expression (right). Traits: RHI - root hair incidence (%), RHL - root hair length (mm), AMF – arbuscular mycorrhizal colonization (%), Arb - arbuscule colonization (%)). Cirsium acaule and Lathyrus pratensis miss RHL measurements, as collected sample material was insufficient for multiple analysis.

**Figure S2:**
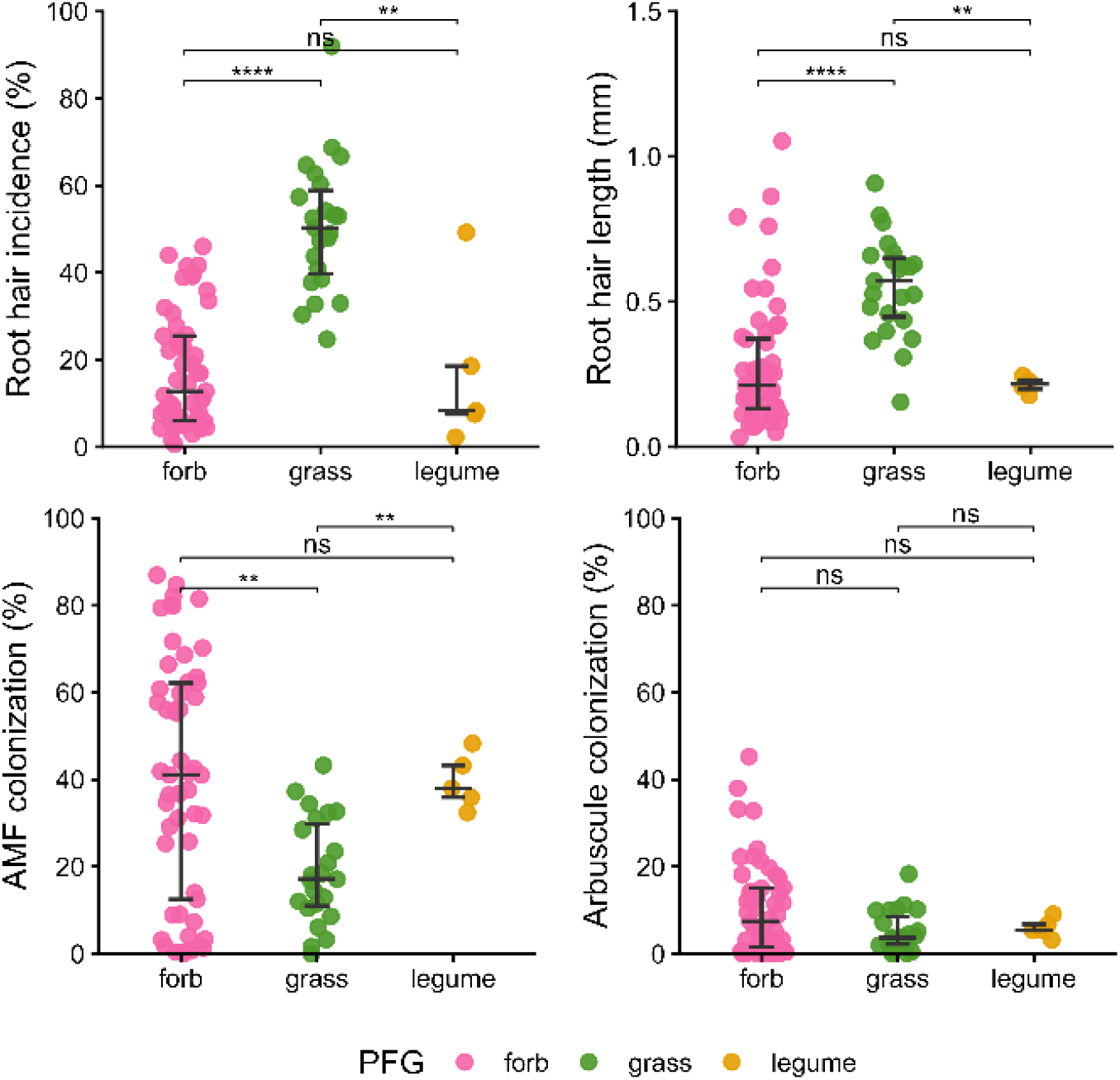
Trait values differ depending on plant functional type (grass, forb, legume), each dot presents a species mean. AMF – arbuscular mycorrhizal fungi, ns – not significant, ** - p-value < 0.01, ** - p-value < 0.001.

**Table S2:** Standardized major axis (SMA) analysis on interspecific scale, split by plant functional type (forb, grass, legume). RHI – root hair incidence, RHL – root hair length, AMF – arbuscular mycorrhizal fungi.

| Trait pair | PFT | slope |
| --- | --- | --- |
| RHI ~ AMF | forb | -0.473 |
| RHI ~ AMF | grass | 1.246 |
| RHI ~ AMF | legume | 3.022 |
| RHI ~ Arbuscule | forb | -1.226 |
| RHI ~ Arbuscule | grass | 3.297 |
| RHI ~ Arbuscule | legume | -8.702 |
| RHL ~ AMF | forb | -0.008 |
| RHL ~ AMF | grass | -0.014 |
| RHL ~ AMF | legume | -0.005 |
| RHL ~ Arbuscule | forb | -0.020 |
| RHL ~ Arbuscule | grass | -0.038 |
| RHL ~ Arbuscule | legume | -0.012 |

**Figure S3:**
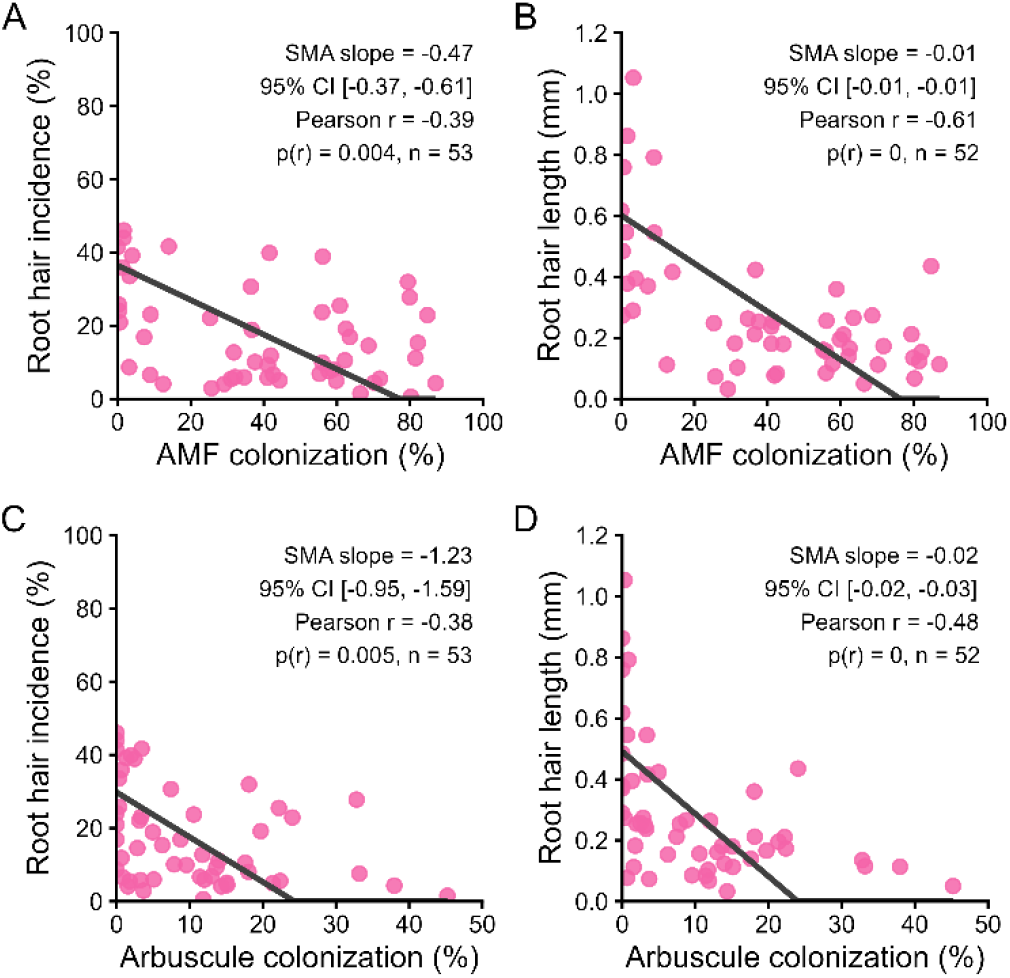
Interspecific relationship of root hair traits and mycorrhizal colonization in the plant functional type of non-leguminous forbs. Species mean trait values of root hair incidence (A, C; n = 53 species) and root hair length (B, D; n = 53 species) were plotted against total AMF colonization (A, B; including all intraradical mycorrhizal structures such as hyphae, arbuscules, vesicles, coils) and arbuscular colonization (C,D). Each point corresponds to a species mean. Black line denotes the standardized major axis (SMA) regression slope. AMF – arbuscular mycorrhizal fungi.

**Figure S4:**
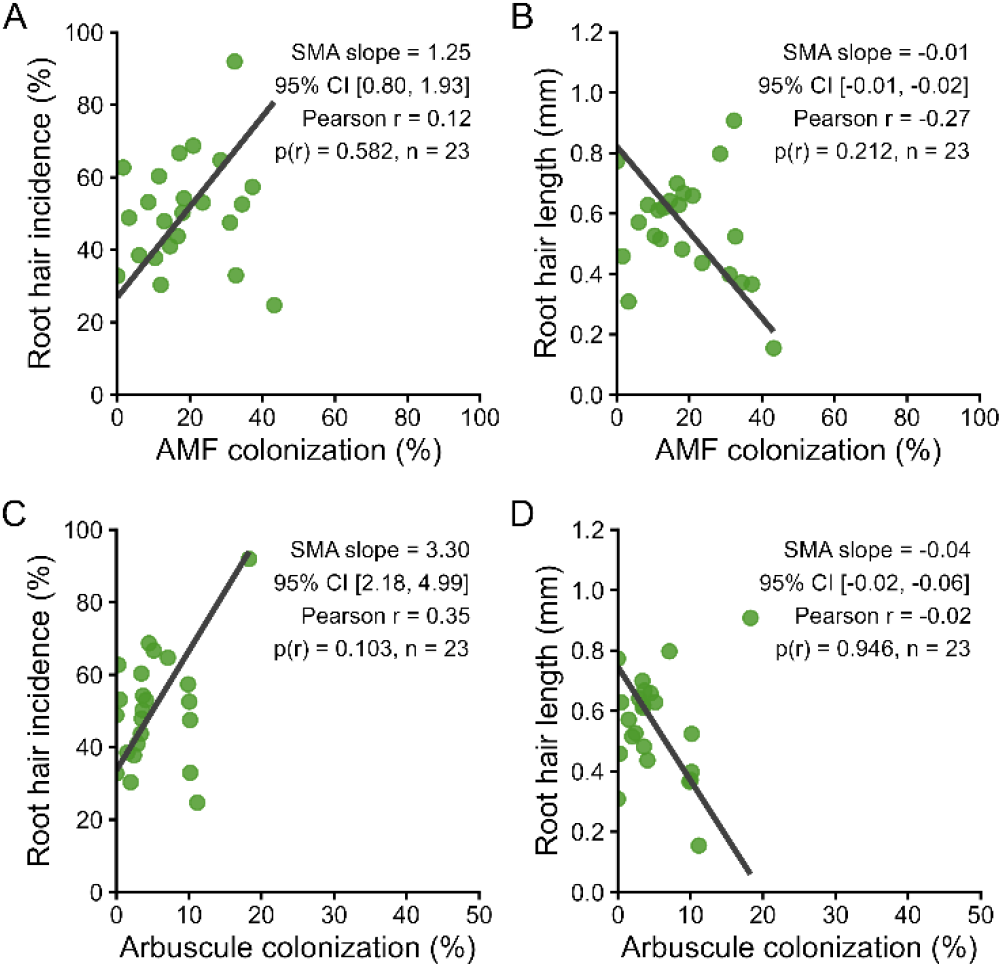
Interspecific relationship of root hair traits and mycorrhizal colonization in the plant functional type of grasses. Species mean trait values of root hair incidence (A, C; n = 23 species) and root hair length (B, D; n = 23 species) were plotted against total AMF colonization (A, B; including all intraradical mycorrhizal structures such as hyphae, arbuscules, vesicles, coils) and arbuscular colonization (C,D). Each point corresponds to a species mean. Black line denotes the standardized major axis (SMA) regression slope. AMF – arbuscular mycorrhizal fungi.

**Figure S5:**
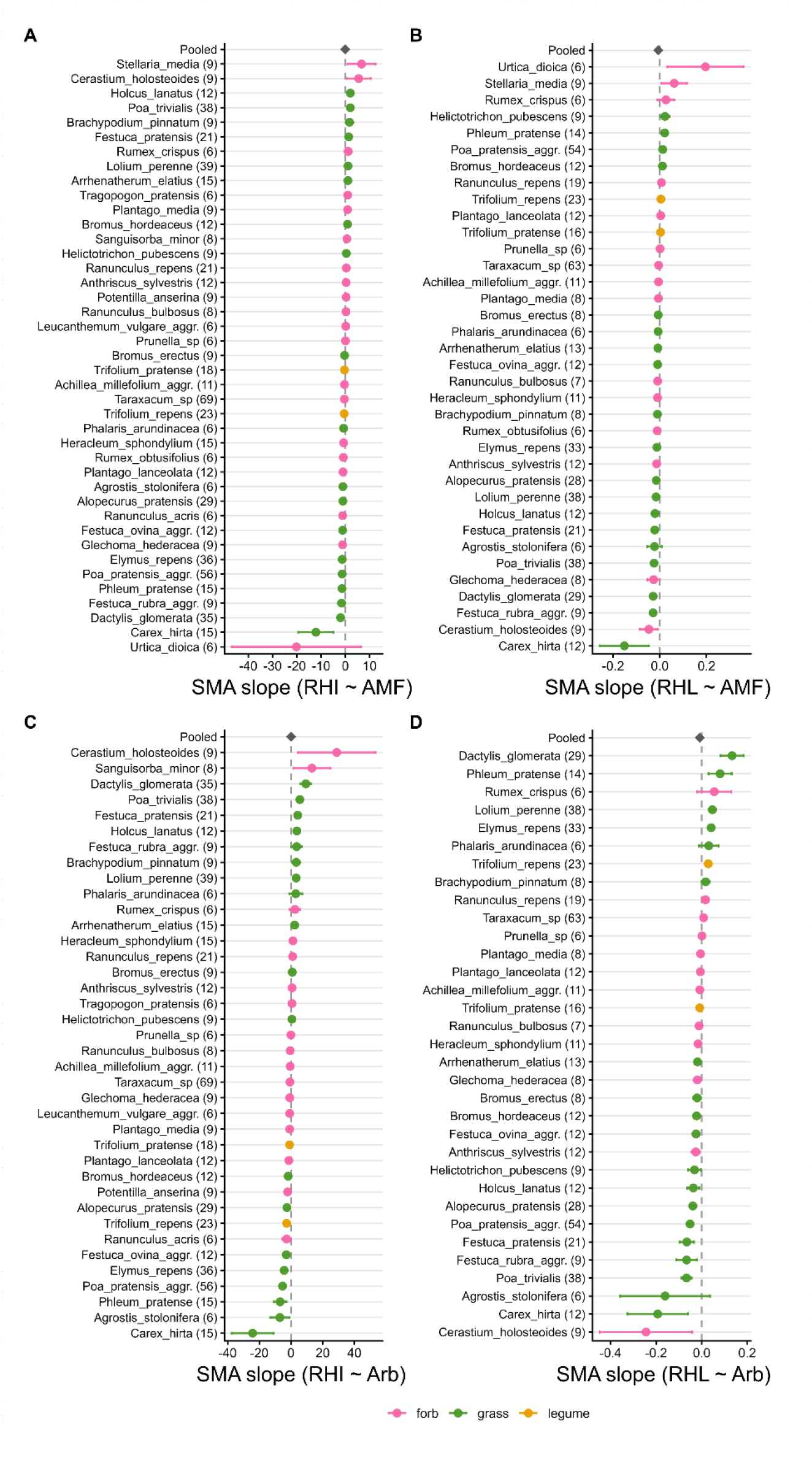
Forest plots showing per-species SMA slope estimate of root hair and mycorrhizal colonization traits, and overall pooled SMA slope estimate (diamoned on top). Number in brackets denotes number of replicates per species used in analysis. Forbs – pink, grasses – green, legumes – amber. RHI – root hair incidence, RHL – root hair length, AMF – arbuscular mycorrhizal colonization, Arb – arbuscule colonization.

**Table S3:** Summary table displaying output of linear models analysing trait-trait relationships on community scale, including the interaction term of grass cover proportion. Reverse models were fitted as a sensitivity analysis with the response and predictor trait reversed; n = 27 plots for all models. RHI – root hair incidence, RHL – root hair length, AMF – arbuscular mycorrhizal fungi colonization (including hyphae, vesicles, arbuscules), Arb – arbuscular mycorrhizal colonization (scoring arbuscules only), CWM – community weighted mean, CI – confidence interval.

| Model type | Response | Predictor | $\beta$ | $p$ | 95% CI lower | 95% CI upper | Adj. R<sup>2</sup> |
| --- | --- | --- | --- | --- | --- | --- | --- |
| Main | RHI CWM | AMF CWM | -0.285 | 0.214 | -0.746 | 0.176 | 0.068 |
| Main | RHL CWM | AMF CWM | -0.361 | 0.0126 | -0.636 | -0.085 | 0.667 |
| Main | RHI CWM | Arb CWM | -0.389 | 0.0779 | -0.826 | 0.047 | 0.086 |
| Main | RHL CWM | Arb CWM | -0.431 | 0.0119 | -0.757 | -0.105 | 0.489 |
| Reverse | AMF CWM | RHI CWM | -0.193 | 0.385 | -0.643 | 0.258 | 0.305 |
| Reverse | AMF CWM | RHL CWM | -0.013 | 0.947 | -0.398 | 0.372 | 0.602 |
| Reverse | Arb CWM | RHI CWM | -0.27 | 0.236 | -0.73 | 0.189 | 0.276 |
| Reverse | Arb CWM | RHL CWM | -0.067 | 0.775 | -0.549 | 0.414 | 0.377 |

**Figure S6:**
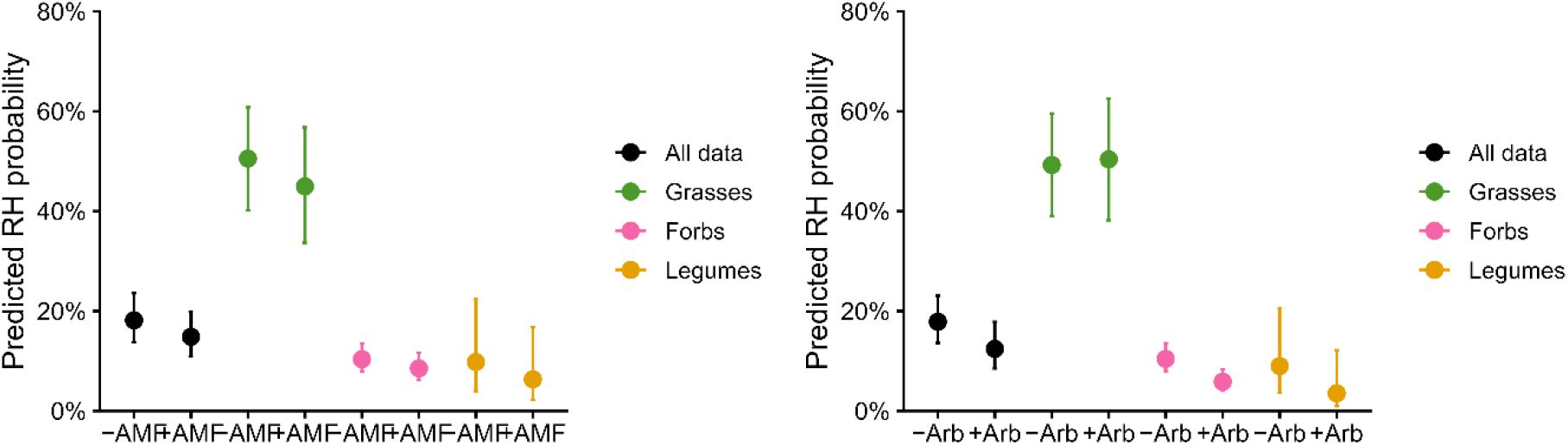
Predicted probabilities of root hair (RH) occurrence depending on AMF (arbuscular mycorrhizal fungi) presence or Arb (arbuscule) presence (y – yes, n – no), across all observations (black) and split by plant functional type.

**Table S4:** Results of logistic mixed-effects models testing associations between the presence of root hairs (RH) and the presence of arbuscular mycorrhizal fungal structures (AMF, including hyphae, vesicles, arbuscules) or arbuscules only (Arb) at the intra-individual scale. Models were fitted in both directions, reversing response and predictor variables. β - estimated regression coefficient on the log-odds scale, SE - standard error, *z* - Wald test statistic, *p* - the associated significance value, OR - odds ratio.

| Model | Response | Predictor | $\beta$ | SE | $z$ | $p$ | OR |
| --- | --- | --- | --- | --- | --- | --- | --- |
| m1 | AMF presence | RH presence | -0.245 | 0.08 | -3.05 | 0.002 | 0.783 |
| m2 | RH presence | AMF presence | -0.241 | 0.08 | -3.134 | 0.002 | 0.786 |
| m3 | Arb presence | RH presence | -0.252 | 0.11 | -2.398 | 0.017 | 0.777 |
| m4 | RH presence | Arb presence | -0.426 | 0.10 | -4.371 | 0.000 | 0.653 |

**Figure S7:**
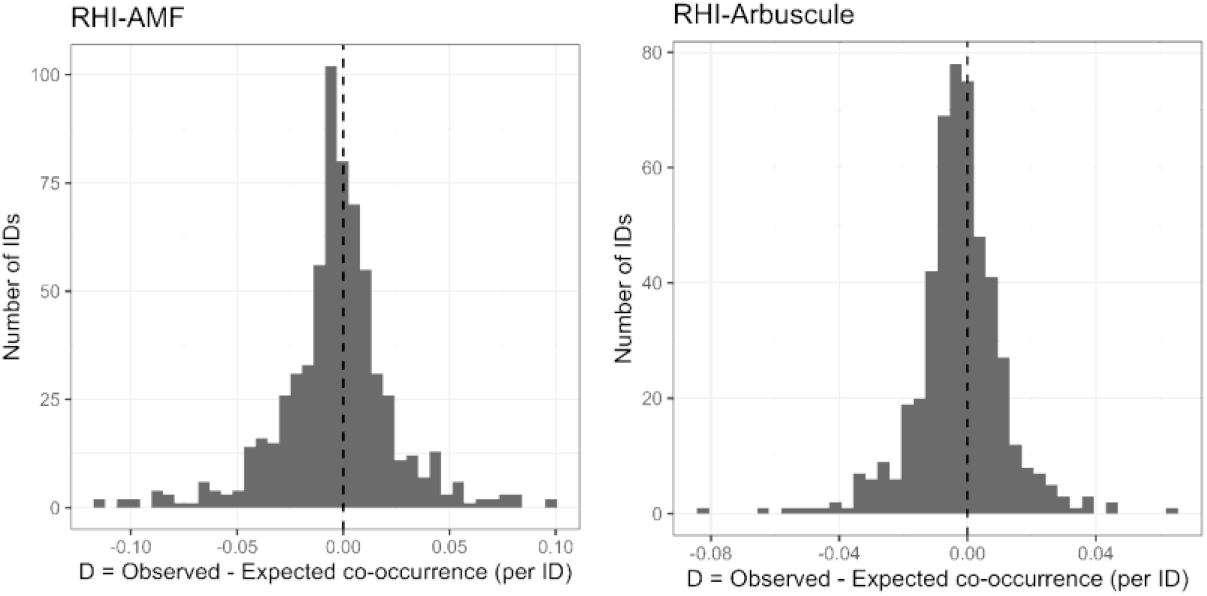
Histogram of deviations (D) between observed and expected co-occurrence of root hairs (RHI) and (A) arbuscular mycorrhizal fungi (AMF) or (B) arbuscules across individuals. Expected co-occurrence was calculated assuming independence of traits. Negative values indicate reduced co-occurrence, positive values indicate increased co-occurrence. The dashed line denotes D = 0. Deviations were evaluated using permutation tests with 2,000 within-individual randomizations.

**Figure S8:**
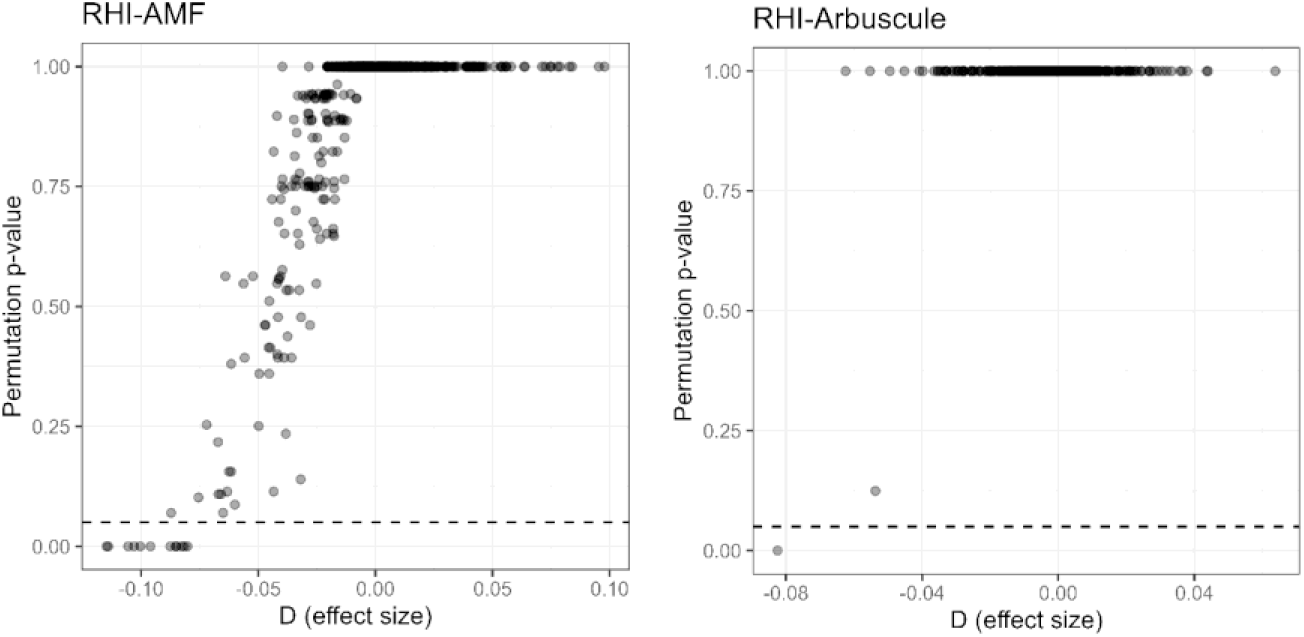
Effect size (D) versus adjusted permutation *p*-values for co-occurrence of root hairs (RHI) and (A) arbuscular mycorrhizal fungi (AMF) or (B) arbuscules across individuals. D is the difference between observed and expected co-occurrence under independence. Negative values indicate reduced co-occurrence. Each point represents one individual. P-values were derived from 2,000 within-individual permutations and corrected for multiple testing (Benjamini–Hochberg). The dashed line marks *p* = 0.05.

